# Transcriptomes resolve phylogenetic relationships and reveal undescribed diversity in taildropper slugs (Genus *Prophysaon*)

**DOI:** 10.64898/2026.03.25.713997

**Authors:** Megan L. Smith, Shelby Moshier, Nathaniel F. Shoobs

## Abstract

The temperate rainforests of the Pacific Northwest of North America harbor many endemic taxa whose evolutionary histories have been shaped by major climatic and geologic events. The enigmatic taildropper slugs (genus *Prophysaon*) are one example, notable for their ability to autonomize their tails to escape predators. Despite extensive work uncovering the evolutionary history of individual lineages, relationships among the nine recognized species of *Prophysaon* remain poorly understood due to insufficient molecular data. To address this, we collected transcriptomes for six of the nine currently accepted species of *Prophysaon*. Using these data, we were able to resolve species relationships, calling into question the existing subgeneric classification based on morphology. We also detected undescribed phenotypic diversity within the *P. andersonii—P. foliolatum* species complex, with molecular data supporting the distinctness of two phenotypically distinct populations from Washington. Finally, our transcriptomic data suggest a moderate role of introgression in shaping the evolutionary history of *Prophysaon*. Here, we synonymize the subgenus *Mimetarion* with nominotypical *Prophysaon*. Future work should further investigate whether the undescribed diversity detected here represents species level differentiation.

## INTRODUCTION

The Pacific Northwest of North America (PNW) is home to two disjunct extents of temperate rainforest—the Cascades and Coastal Ranges in the west and the Northern Rocky Mountains in the east. These rainforests are home to more than 150 rainforest endemics (Nielson *et al*., 2001) whose evolutionary histories and current distributions have been shaped by major geologic and climatic events. The two extents of rainforests are separated by over 200 km by the arid shrub-steppe habitat of the Columbia Basin, which formed due to the orogeny of the Cascades Range around 2-5 Mya (Graham, 1999). Although the Central Oregonian Highlands and the Okanogan Highlands lessen the isolation between inland and coastal rainforests, the Columbia Basin has served as a major barrier to dispersal for many rainforest endemics (e.g., Nielson *et al*., 2001, 2006; Carstens *et al*., 2004, 2005b,a; Steele *et al*., 2005). Furthermore, Pleistocene climatic fluctuations had major impacts on the distributions of rainforest endemics. During the height of glaciation, glaciers covered large portions of species’ current ranges, which may have eliminated species from northern portions of their ranges or forced them into isolated refugia (Brunsfeld *et al*., 2001; Carstens *et al*., 2005a; Smith *et al*., 2022b), potentially leading to secondary contact and introgression between previously isolated populations and species.

*Prophysaon* Bland and Binney, 1874 [“1873”] is an enigmatic genus of terrestrial gastropods endemic to the temperate rainforests of the PNW, ranging from Alaska to northern California. Commonly known as the taildropper slugs, all *Prophysaon* species are capable of caudal autotomy when threatened (Pilsbry, 1948). The taxonomy of the genus is in need of revision due to the overlapping distributions and phenotypes of many of its members and poor circumscription of the available names and constituent taxa (Wilke & Duncan, 2004; Smith *et al*., 2018). The 9 currently accepted species in the genus (Supporting Fig. S1) are divided into two subgenera: nominotypical *Prophysaon* Bland and Binney, 1874 and *Mimetarion* Pilsbry, 1948. Pilsbry described the subgenus *Mimetarion* with *P. vanattae* Pilsbry, 1948 as its type, and *P. fasciatum* (Cockerell in W.G. Binney, 1890) and *P. humile* Cockerell, 1890 as members, and placed *P. andersonii* (Cooper, 1872)*, P. boreale* Pilsbry, 1948*, P. coeruleum* Cockerell, 1890*, P. dubium* (Cockerell, 1890), and *P. foliolatum* (Gould in A. Binney, 1851) in the nominotypical subgenus *Prophysaon*, on the basis of genital morphology differences between these two groups of species (1948). The species of *Mimetarion* lack the specialized massive and convoluted epiphallus typical of *P. (Prophysaon)* (Pilsbry, 1948).

*P. andersonii* is the most widespread of the taildropper slugs, occurring throughout the coastal and Cascades ranges and the Northern Rockies in large numbers. *P. foliolatum* is difficult to distinguish from *P. andersonii* phenotypically and is distributed in western Washington. *P. coeruleum* is a distinct species with blue-grey coloration occurring in both the Cascades and the Northern Rocky Mountains. Disjunct populations of *P. dubium* are found in the Cascades and Coastal ranges, northern Idaho, and northeastern Washington (Burke, 2013). *P. vanattae* occurs only in the Cascades mountains, while *P. humile* occurs only in the Northern Rocky Mountains, suggesting that the orogeny of the Cascades and subsequent aridification of the Columbia Basin may have played a role in their divergence. *P. obscurum* Cockerell sensu Branson (1977) is a questionable taxon that has a restricted geographic range, occurring south of South Puget Sound and extending through western Washington and the Columbia Gorge (Burke, 2013). Limited molecular data suggest that *P. obscurum* Cockerell sensu Branson may be conspecific with *P. vanattae* (Wilke & Duncan, 2004; Smith *et al*., 2018), though without examining Cockerell’s type specimens and sequencing topotypic specimens, it is unclear whether *obscurum* is the correct name for the populations that Branson (1977) and later authors have collected and sequenced.

Recent work has investigated population structure within species of *Prophysaon*, focusing primarily on those species with disjunct distributions across the Columbia Basin and sister-species pairs distributed across the Columbia Basin. Smith et al. (2018) used data from the mitochondrial gene COI to test for deep divergence across the Columbia Basin associated with the orogeny of the Cascades range, but found that the data best supported recent dispersal to the inland rainforests. Despite the lack of intraspecific divergence across the Columbia Basin, the results suggested deeper population structure in the coastal and Cascades ranges. Subsequent work used reduced representation genomic sequencing (Genotyping-by-Sequencing; Elshire *et al*., 2011) to investigate the role of Pleistocene glacial cycles in shaping genetic variation in *Prophysaon* in the coastal and Cascades ranges. This work supported divergences dating to the late Pleistocene and secondary contact between lineages, suggesting a potential role of Pleistocene glacial cycles in shaping intraspecific genetic variation in *Prophysaon* slugs.

Despite this recent progress, relationships among species remain poorly understood. The gene tree inferred from COI data failed to resolve relationships among species (Smith *et al*., 2018), and GBS data were not suitable for phylogenetic reconstruction due to limited loci shared across some species. Here, we sequenced transcriptomes from 6 of the 9 species of *Prophysaon* recognized by Turgeon et al. (1998). We were able to resolve relationships among species, and our results further suggest that the current morphology-based subgeneric classification is of limited utility. We also uncovered potentially undescribed diversity in the genus. Finally, our data support a moderate role of introgression in the history of *Prophysaon*.

## MATERIALS AND METHODS

### Sampling and sequencing

We collected slug specimens from the Pacific Northwest during the autumns of 2020 and 2021. In 2020, we placed slugs directly into RNALater upon collection. In 2021, we dissected slugs first to improve tissue penetration with RNA Later. Total RNA was extracted using the TRIzol reagent (Thermo Fisher Scientific) following the manufacturer’s protocol. RNA was extracted either from several tissues (albumin gland, digestive gland, epiphallus, mantle covering, and foot), from all of those tissues except the digestive gland, from only the albumin gland, or from the albumin gland and the epiphallus (Supporting Table S1). While initial extractions focused on using several tissues, for samples with poor quality extractions, using only the albumin gland and epiphallus improved quality. Because of this, we focused on only the albumin gland and epiphallus for all extractions from slugs collected in 2021. An Agilent TapeStation 4200 was used to assess total RNA quality and determine suitability for sequencing. Illumina TruSeq Stranded mRNA HT Libraries were prepared by the Center for Genomics and Bioinformatics at Indiana University and sequenced on an Illumina NextSeq500 system using 150 bp paired-end sequencing.

### Transcriptomic data—processing and assembly

All transcriptomes were assembled *de novo* following the pipeline used in Cunha and Giribet (2019) and associated scripts. Raw reads were error-corrected using RCorrector v1.0.4 (Song & Florea, 2015), and unfixable reads were removed. TrimGalore! v0.6.0 (Krueger, 2019) was used to remove adapters and reads shorter than 50 bp. We then assessed read quality using FastQC v0.11.9 (Andrews, 2010). We compared the filtered reads to a databased of rRNAs and removed matching reads using Bowtie2 v2.4.2 (Langmead & Salzberg, 2012). This database was created from an 18S sequence from the slug *Arion ater* from the SILVA database (Accession: HQ659992) and a 5S sequence from *Arion rufus* from GenBank (Accession: J01888). Next, filtered reads were assembled into transcripts using Trinity v2.11.0 (Haas *et al*., 2013). We reran Bowtie2 on assembled transcripts to remove any remaining rRNA from assemblies and used CD-HIT-EST v4.6.8 (Fu *et al*., 2012) to remove redundant transcripts (sequence identity > 95%).

Transcripts were translated to amino acids using TransDecoder v5.5.0 (Haas *et al*., 2013), and the longest isoform was retained using a custom script from Cunha and Giribet (2019). For use as an outgroup in phylogenetic analyses, we downloaded raw RNA-seq data from *Arion vulgaris* from GenBank (Accession: SRX9728973), and we followed the same steps as above to process reads and assemble the transcriptome. The completeness of all assemblies was assessed using BUSCO v5.0.0 (Manni *et al*., 2021) and the Mollusca database.

### Phylogenetic inference

To infer relationships among species, we assembled phylogenomic datasets using a subset of the sequenced transcriptomes. We included the outgroup, *Arion vulgaris*, and selected the two most complete transcriptomes per species when possible. For *P. humile* and *P. coeruleum* only a single transcriptome was available. We also included two samples that appeared to be members of the *Prophysaon andersonii* species complex (consisting of *P. andersonii* and *P. foliolatum*), but which were morphologically distinct from typical specimens of *P. andersonii* and *P. foliolatum* (see Results). In total, we included 13 transcriptomes for our phylogenomic analyses. To identify ortholog clusters, we first performed an all-by-all BLASTN search on the CDS of all 13 specimens using blast v2.9.0 (Camacho *et al*., 2009). Then, we used a custom script from Yang and Smith (2014) to concatenate blast results and filter results with a hit fraction less than 0.4. The mcl algorithm (van Dongen, 2000) with an inflation parameter of 1.4 and a hit fraction cutoff of 0.3 was used to cluster the filtered BLASTN output. Sequences were aligned by codon using GUIDANCE2 v2.0.2 (Sela *et al*., 2015) with MAFFT v7.490 (Katoh & Standley, 2013) and 100 bootstrap replicates. We used trimAI v1.2rev59 (Capella-Gutiérrez *et al*., 2009) to remove all positions in the alignment with gaps in 50% or more of the sequences. Finally, we filtered alignments shorter than 200 bp, removed individuals whose sequences were composed of more than 50% gaps from each alignment, and removed alignments with fewer than four sequences.

We inferred a gene tree for each orthogroup using IQTree v2.0.6 (Minh *et al*., 2020) with the best model of nucleotide substitution selected using ModelFinder (Kalyaanamoorthy *et al*., 2017), as implemented in IQTree. From these gene trees, we generated several datasets for downstream species tree inference. Phylogenetic inference has typically relied on single-copy gene families, as these families are likely to contain only orthologs, or genes related through speciation events, to the exclusion of paralogs, or genes related through duplication events.

However, recent work has demonstrated that phylogenetic inference using large gene families, including paralogs, is accurate (Zhang *et al*., 2020; Smith & Hahn, 2021; Yan *et al*., 2022; Smith *et al*., 2022a). We created a series of datasets differing in their inclusion of data from larger gene families. First, we analyzed only single-copy gene families. Then, we used three branch-cutting approaches to extract putative single-copy genes from larger gene families: Maximum Inclusion and Monophyletic Outgroups (Yang & Smith, 2014) and DISCO (Willson *et al*., 2022). Finally, we created a dataset of all gene families, including both orthologs and paralogs.

Next, we analyzed these datasets using several approaches that vary in their ability to accommodate discordance due to incomplete lineage sorting (ILS) and gene duplication and loss (GDL). First, all datasets were analyzed using ASTRAL-III v5.7.3 (Zhang *et al*., 2018), which performs well in the presence of GDL and ILS (Legried *et al*., 2021) We also analyzed the dataset including all gene families using ASTRAL-Pro v1.1.5 (Zhang *et al*., 2020), which was designed specifically for analyzing gene families including paralogs. Finally, we generated concatenated datasets for the single-copy genes and all datasets generated using branch-cutting approaches and inferred species trees from these concatenated datasets using IQTree. The best model of nucleotide substitution was inferred using ModelFinder, as above, and 1000 ultrafast bootstrap replicates were used to assess nodal support. To assess the amount of discordance in these datasets, we calculated both gene concordance factors (GCFs) and site concordance factors (SCFs) in IQ-Tree (Minh *et al*., 2020). As a reference tree, we used the ASTRAL tree inferred from single-copy orthologs, and 100 quartets were used to estimate SCFs.

### Inferring population structure and testing for introgression

Next, we applied population genetic approaches to investigate the evolutionary history of the *P. andersonii—P. foliolatum* species complex, given the presence of potential undescribed diversity in this group. For these analyses, we mapped filtered reads against the *Arion vulgaris* genome, as this is the closest-available reference for the focal taxa. The reference genome was downloaded from NCBI (Accession: ASM2079622v1) and indexed using bwa v.0.7.12 (Li, 2013). Filtered reads were then mapped to the reference using bwa-mem v2.0pre2 (Li, 2013). Samtools v1.15.1 (Li *et al*., 2009) fixmate was used to fill in read-pairing info and convert to .bam format. Then, samtools markupd was used to mark and remove duplicates. Reads with low quality mapping (q < 20) were removed using samtools, and overlapping paired-end reads were removed using bamutil’s clipOverlap function (Jun *et al*., 2015).

Next, filtered reads were used to call genotypes using bcftools v1.22 (Li, 2011), excluding low quality base calls (q < 20) and indels. Then, we used vcftools v0.1.17 (Danecek *et al*., 2011) to filter genotypes. We removed invariant sites, sites with more than two alleles, sites with low-quality genotypes (q < 20), and sites with extreme depth values (mean depth < 3 or > 60). We also removed rare variants (maf <=5%), and sites with high levels (>50%) of missing data. For Structure analyses (see below), we included only individuals from the *P. andersonii—P. foliolatum* complex and retained only a single SNP per 5000 bp window (--thin 5000) to minimize the effects of linkage. To investigate introgression in Dsuite (see below), we created a second dataset including all individuals and no thinning.

To investigate population structure within this species complex, we used the program Structure v2.3.4 (Pritchard *et al*., 2000) to analyze the dataset from the *P. andersonii—P. foliolatum* complex. We used PGDSpider v2.1.1.5 (Lischer & Excoffier, 2012) to convert the filtered vcf into structure-formatted input. Structure analyses were run with k-values from one to seven with ten replicates per k. The first 100,000 generations were discarded as burn-in, followed by 500,000 samples. We used pophelper (Francis, 2017) to summarize results, implement the Evanno method for choosing k (Evanno *et al*., 2005), and visualize results.

To investigate signatures of introgression, we used Dsuite v0.5 (Malinsky *et al*., 2021) and the dataset with all individuals. We calculated D and f4-ratio statistics for all possible trios and calculated f-branch statistics using the ASTRAL species tree as the guide tree and *Arion vulgaris* as the outgroup.

### Mitochondrial data

To place the newly discovered diversity within the *P. andersonii–P. foliolatum* complex into a broader phylogenetic context, we extracted the mitochondrial gene COI from our RNAseq data and compared it to all COI data available from *Prophysaon* on GenBank. We extracted COI from raw reads using MitoGeneExtractor (Brasseur *et al*., 2023), which uses Exonerate to construct alignments (Slater & Birney, 2005). As the protein reference, we used GenBank accession AUH20884 from *P. andersonii.* We used the invertebrate mitochondrial code and the default values for all other settings. We aligned the extracted sequences and *Prophysaon* COI from GenBank using muscle v3.8.31 (Edgar, 2004). Then, we selected the best model of nucleotide substitution, reconstructed the maximum likelihood tree, and assessed support using the ultrafast bootstrap in IQTree v2.3.2 (Minh *et al*., 2020).

### Morphological data

Radulae (n=39) were dissected from a subset of species and specimens collected for this and previous studies (Smith *et al*., 2018, 2022b), cleaned in a potassium hydroxide solution as needed, and mounted on carbon tape for scanning electron microscopy. Radulae were imaged on SEMs at either the Center for Electron Microscopy and Analysis at The Ohio State University, or Indiana University’s Electron Microscopy Center. To better examine the range of variation in radular morphology within and between species, we focused on the *P. andersonii—P. foliolatum* species complex, given better sampling within this complex. We used geometric morphometrics to analyze shape variation in the rachidian (central) teeth by species, focusing on a single exemplar tooth per specimen. We placed 6 fixed landmarks and computed 4 curves (Supporting Fig. S2) using the R package stereomorph (Olsen & Westneat, 2015). We then used a Generalized Procrustes Analysis as implemented in the gpagen function of the R package geomorph (Adams & Otárola-Castillo, 2013).

To visually assess differences between populations in shape, we performed a PCA using the gm.prcomp function in the geomorph package. To test whether variation in shape is explained by population, we performed a permutational Procrustes ANOVA using the function procD.lm in geomorph, with 10,000 permutations. We used Type 1 Sums of Squares and included centroid size as the first covariate. Then, to determine which populations differed in shape, we used the pairwise function in the R package RRPP (Collyer & Adams, 2018) and corrected for multiple comparisons using a Bonferroni correction.

## RESULTS

### Sampling, Sequencing, and Assembly

We collected representatives (Fig. 1) of *Prophysaon andersonii, P. coeruleum, P. dubium, P. foliolatum, P. humile, and P. vanattae*, 6 of the 9 *Prophysaon* species treated as valid by Turgeon et al. (1998). All species were sampled from the western portion of their range, except for *P. humile*, which only occurs inland in the Northern Rocky Mountains of Idaho and Montana. We were unable to collect specimens of the nominal species *P. boreale*, *P. fasciatum*, and *P. obscurum,* all of which we consider nomina dubia (see Discussion). Though we did not focus on collecting topotypic material for this study, we indicate the type locality of all 9 *Prophysaon* species in the distribution maps (Supporting Fig. S1).

**Figure 1.**
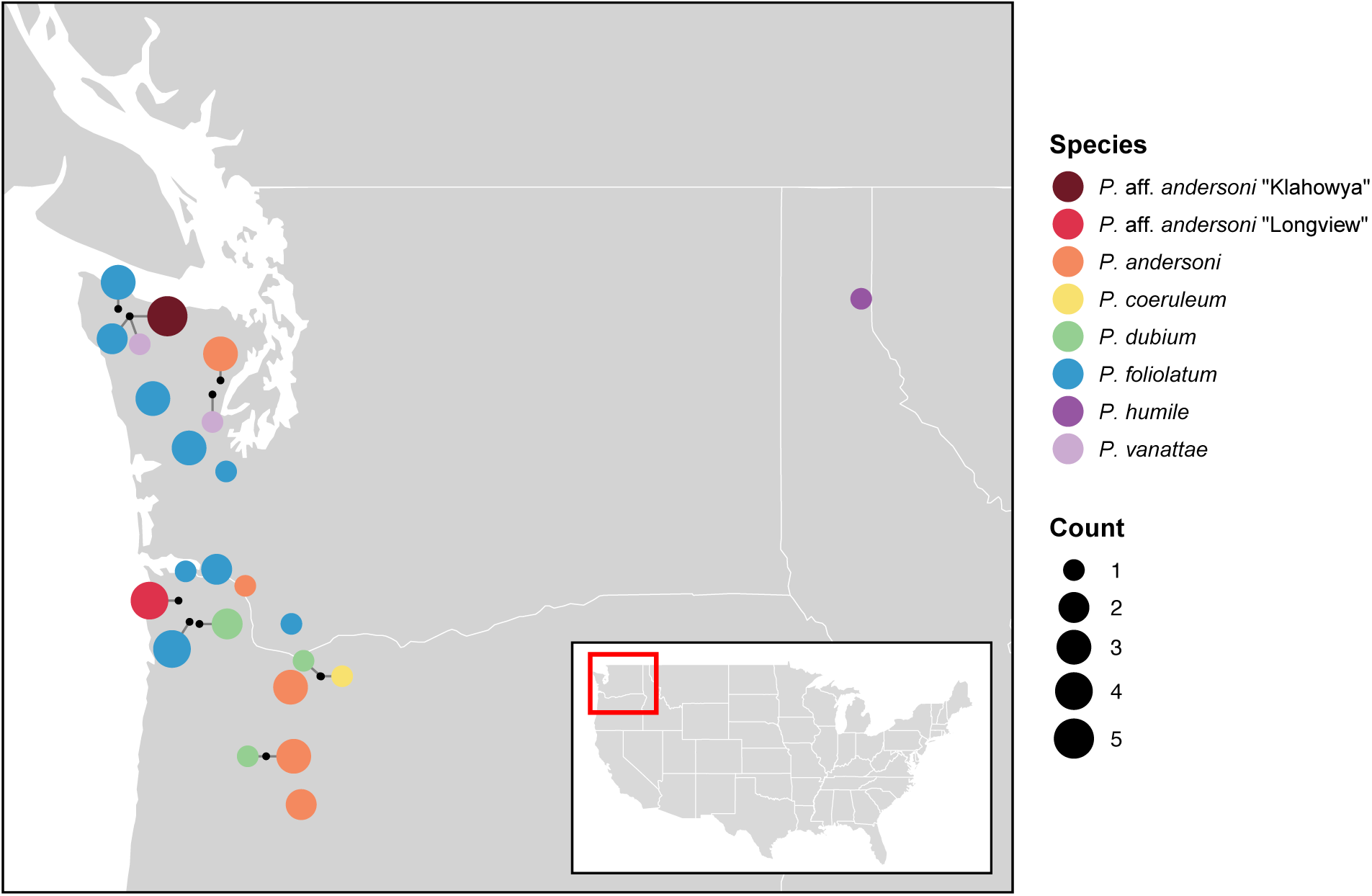
Sampling map of *Prophysaon* specimens collected for this study. The size of points indicates the number of slugs collected from each species at each locality.

In addition to the 6 named species sampled, we found potentially undescribed phenotypic and genetic diversity within the *P. andersonii—P. foliolatum* complex. Some specimens from near Longview, Washington (*P.* aff. *andersonii* “Longview”, hereafter) appeared similar to *P. foliolatum* and *P. andersonii,* except that they had solid dark brown coloration on the body and mottled patterning on the mantle (Fig. 2; Supporting Fig. S3). Specimens from the Klahowya Campground in Olympic National Forest (*P.* aff. *andersonii* “Klahowya”, hereafter) were also similar to *P. foliolatum* and *P. andersonii* except that they had mottled brown patterning (Fig. 2; Supporting Fig. S3).

**Figure 2.**
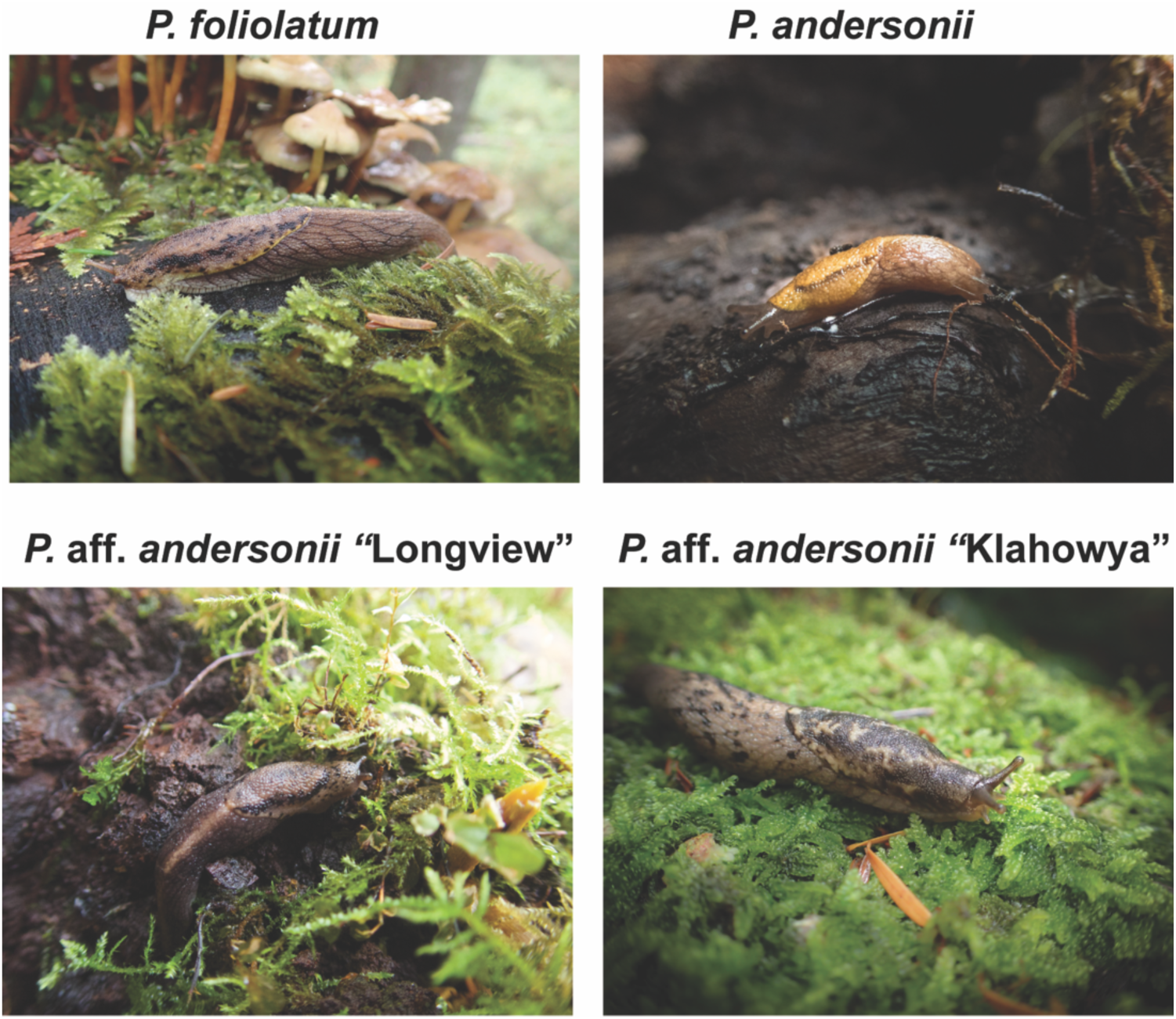
Images of *P. andersonii, P. foliolatum*, and phenotypically unique slugs from Longview, Washington and the Klahowya campground in Olympic National Forest. All photos taken by M. Smith.

We assembled between 23,222 and 128,830 genes using Trinity (Supporting Table S1). The fraction of complete BUSCO genes assembled range from 10 to 55 percent (Supporting Table S1). While these percentages are relatively low, we were able to assemble a sufficient number of genes for phylogenetic and population genetic inference (see below).

### Phylogenetic inference

The dataset consisting of only single-copy orthologs included 7,315 gene families, while the dataset consisting of all gene families (i.e., orthologs and paralogs) included 8,081 gene families (Supporting Table S3). The topology was consistent across all datasets and inference methods (Fig. 3, Supporting Fig. S4). In both ASTRAL and ASTRAL -Pro, local posterior probabilities were 1.0 across all branches and datasets, except ASTRAL-DISCO, where two branches had local posterior probabilities > 0.999. Bootstrap support values were also 100 across all branches and datasets.

**Figure 3.**
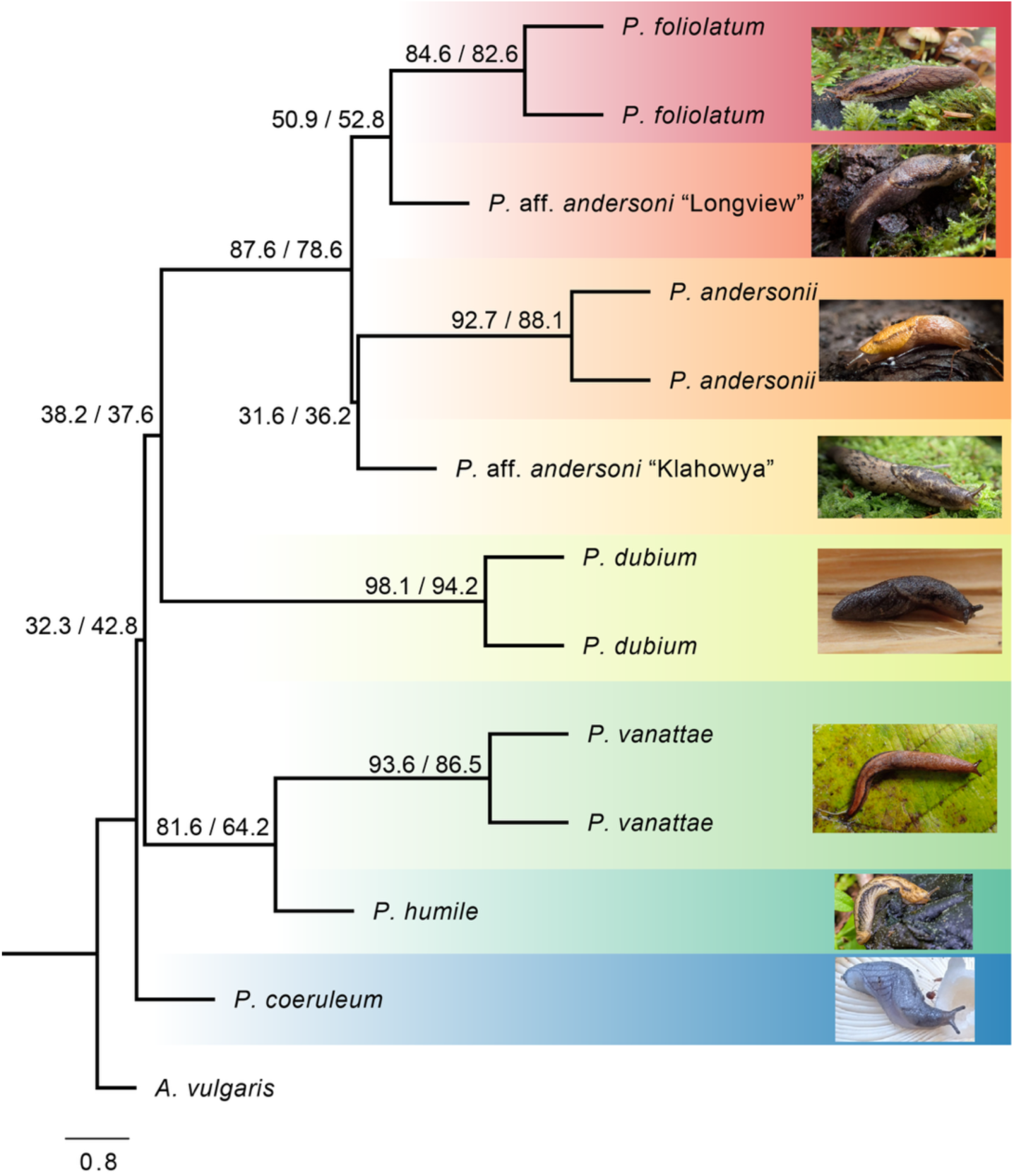
Species tree inferred from single-copy gene trees using ASTRAL. Tree topologies were identical across ASTRAL and ASTRAL -Pro for all datasets analyzed. Node labels show GCFs/SCFs for the dataset including only single-copy gene families. Local posterior probabilities in ASTRAL and ASTRAL -Pro were 1.0 across all nodes and datasets, except ASTRAL-DISCO where two nodes had local posterior probabilities > 0.999. Photos of *P. foliolatum*, *P.* aff. *andersonii* “Longview”, *P. andersonii*, *P.* aff. *andersonii* “Klahowya”, *P. dubium*, and *P. vanattae* by M. Smith. *P. humile* [photo](https://www.inaturalist.org/observations/120327664) by iNaturalist user Radd Icenoggle used under [CC-BY-NC 4.0](https://creativecommons.org/licenses/by-nc/4.0/). *P. coeruleum* [photo](https://www.inaturalist.org/observations/120327664) by iNaturalist user tomcat_clover used under [CC-BY-NC 4.0](https://creativecommons.org/licenses/by-nc/4.0/).

We recovered the *P. andersonii*—*P. foliolatum* species complex with high gene and site concordance factors. Within this species complex, *P.* aff. *andersonii* “Longview” and *P.* aff. *andersonii* “Klahowya” were not recovered as sister to each other. Instead, *P. andersonii* was sister to *P.* aff. *andersonii* “Klahowya”, and *P. foliolatum* was sister to *P.* aff. *andersonii* “Longview”. Gene and site concordance factors were very low for the branch uniting *P. andersonii* and *P.* aff. *andersonii* “Klahowya”, suggesting high ILS and short speciation times.

Relationships between species did not support the current subgeneric level taxonomy. *P. dubium* was recovered as sister to the *P. andersonii—P. foliolatum* species complex, but *P. coeruleum* was recovered as sister to all other *Prophysaon,* including a clade containing *P. vanattae* and *P. humile*, which are currently a part of the subgenus *Mimetarion*, making subgenus *Prophysaon* paraphyletic. Notably, site and gene concordance factors were relatively low in the backbone of the tree, particularly for the branch placing *P. coeruleum* as sister to all other *Prophysaon* and the branch uniting *P. dubium* and the *P. andersonii—P. foliolatum* species complex.

### Inferring population structure and testing for introgression

The dataset used for structure (i.e., with thinning, and only including individuals from the *P. andersonii—P. foliolatum* species complex) included 3,650 variants. While the ΔK method supported K=2, Ln(K) continued to increase up to K=4 (Supporting Fig. S5). At K=2, Structure identified two populations: *P. andersonii*, and a population consisting of *P. foliolatum* and *P.* aff. *andersonii* “Klahowya” and “Longview”. However, *P.* aff. *andersonii* “Klahowya” appeared admixed. At K=3, *P.* aff. *andersonii* “Klahowya” was also identified a distinct population, and *P.* aff. *andersonii* “Longview” appeared as admixed between this population and *P. foliolatum*.

At K=4, *P. andersonii, P. foliolatum*, *P.* aff. *andersonii* “Klahowya”, and *P.* aff. *andersonii* “Longview” were all distinct, with minimal evidence of potential admixture (Fig. 4). While ΔK peaked again at K=6, there was no evidence of additional structure at K-values higher than 4 (Supporting Fig. S6).

**Figure 4.**
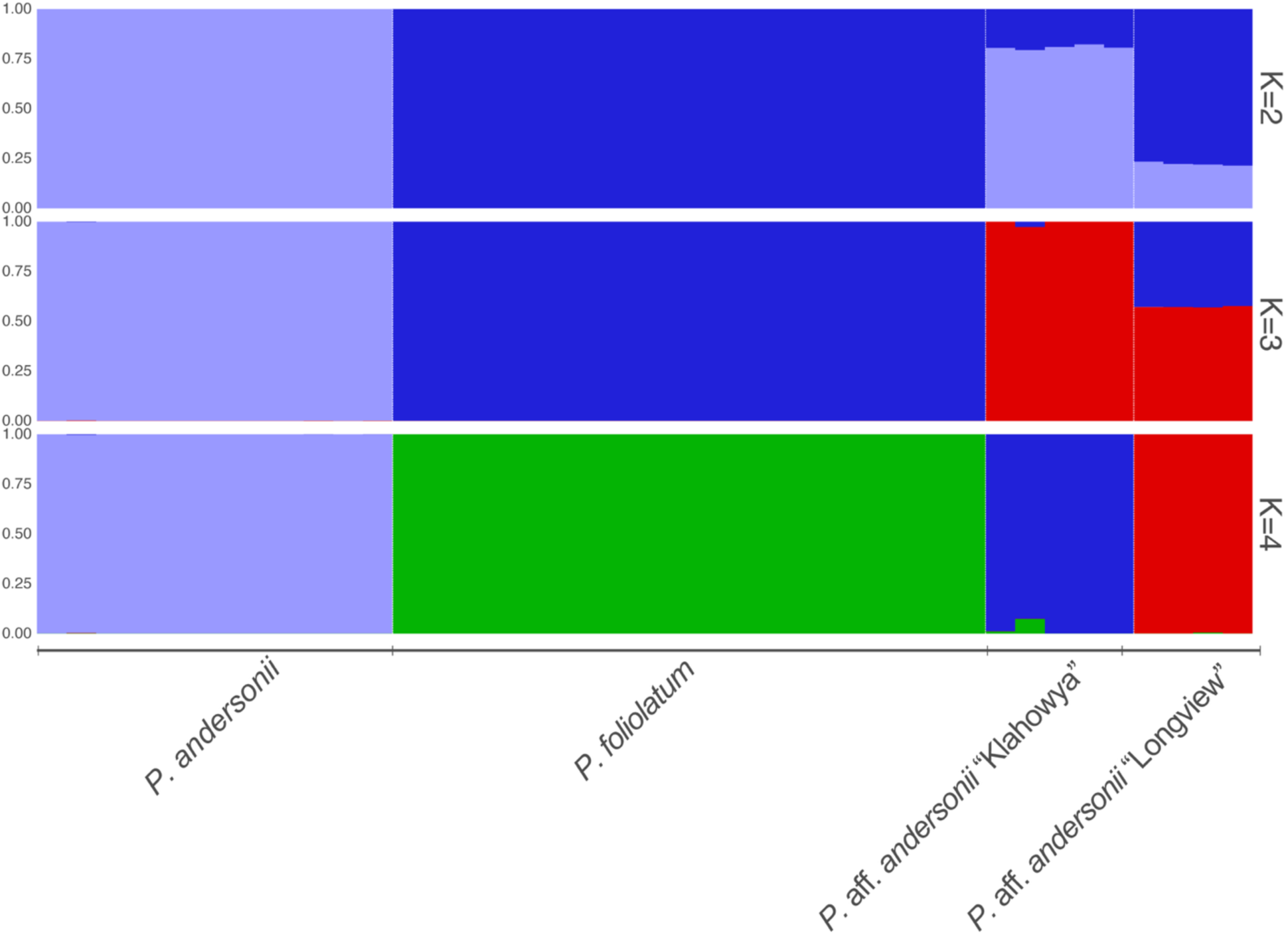
Structure results for K=2 through K=4. Colors indicate cluster assignment probabilities for each individual.

**Figure 5.**
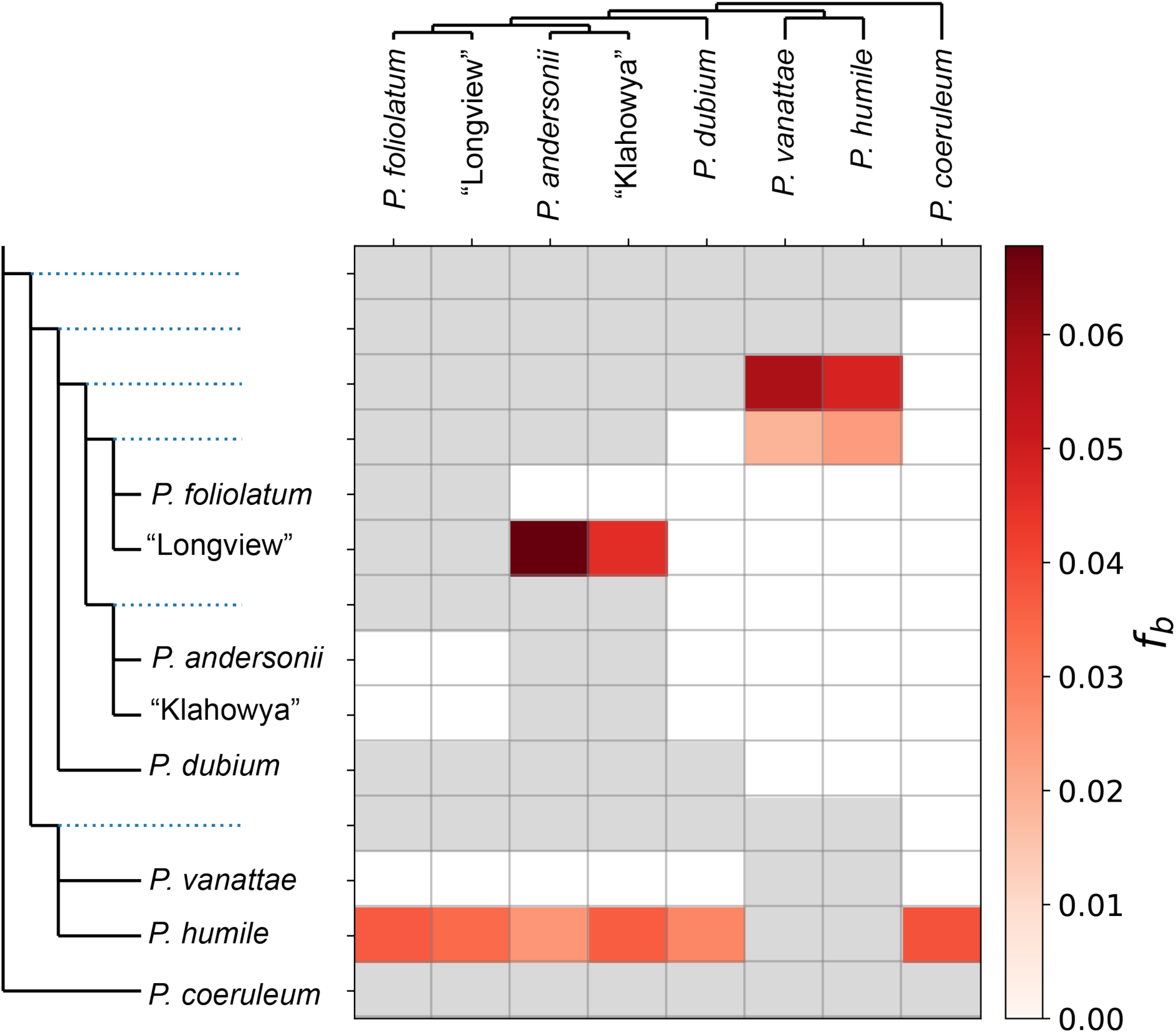
Patterns of introgression based on the *f-*branch test. The color gradient represents the *fb* score, which indicates the degree of excess allele sharing. Greyed out boxes indicate tests that are inconsistent with the species tree. Colored boxes indicate results that are significant at an α of 0.01.

The dataset used for Dsuite (i.e., without thinning, and including all individuals) included 35,512 variants. When D-statistics were calculated based on the user-supplied species tree, 26.8% of tests were significant, suggesting a moderate role of introgression in the history of *Prophysaon*. The *f*-branch statistic identified excess allele sharing between *P.* aff. *andersonii* “Longview” and *P. andersonii* and *P.* aff. *andersonii* “Klahowya”. There was also evidence of potential introgression between *P. vanattae* and *P. humile* and the *P. andersonii*—*P. foliolatum* species complex. Finally, we found evidence of excess allele sharing between *P. humile* and all clades for which valid tests could be performed.

### Mitochondrial data

To investigate the relationships of the phenotypically distinct populations to other populations of *Prophysaon* more broadly, we extracted the mitochondrial gene COI from our transcriptomes and aligned it with COI from GenBank. As expected based on previous studies (e.g., Wilke & Duncan, 2004; Smith *et al*., 2018), relationships among species were not well resolved, with very low bootstrap support (Fig. 6; Supporting Figs. S7-S8). In contrast to the findings from the transcriptomes, *P.* aff. *andersonii* “Klahowya” and “Longview” were recovered as sister in the mitochondrial tree, and they were nested within *P. andersonii*.

**Figure 6.**
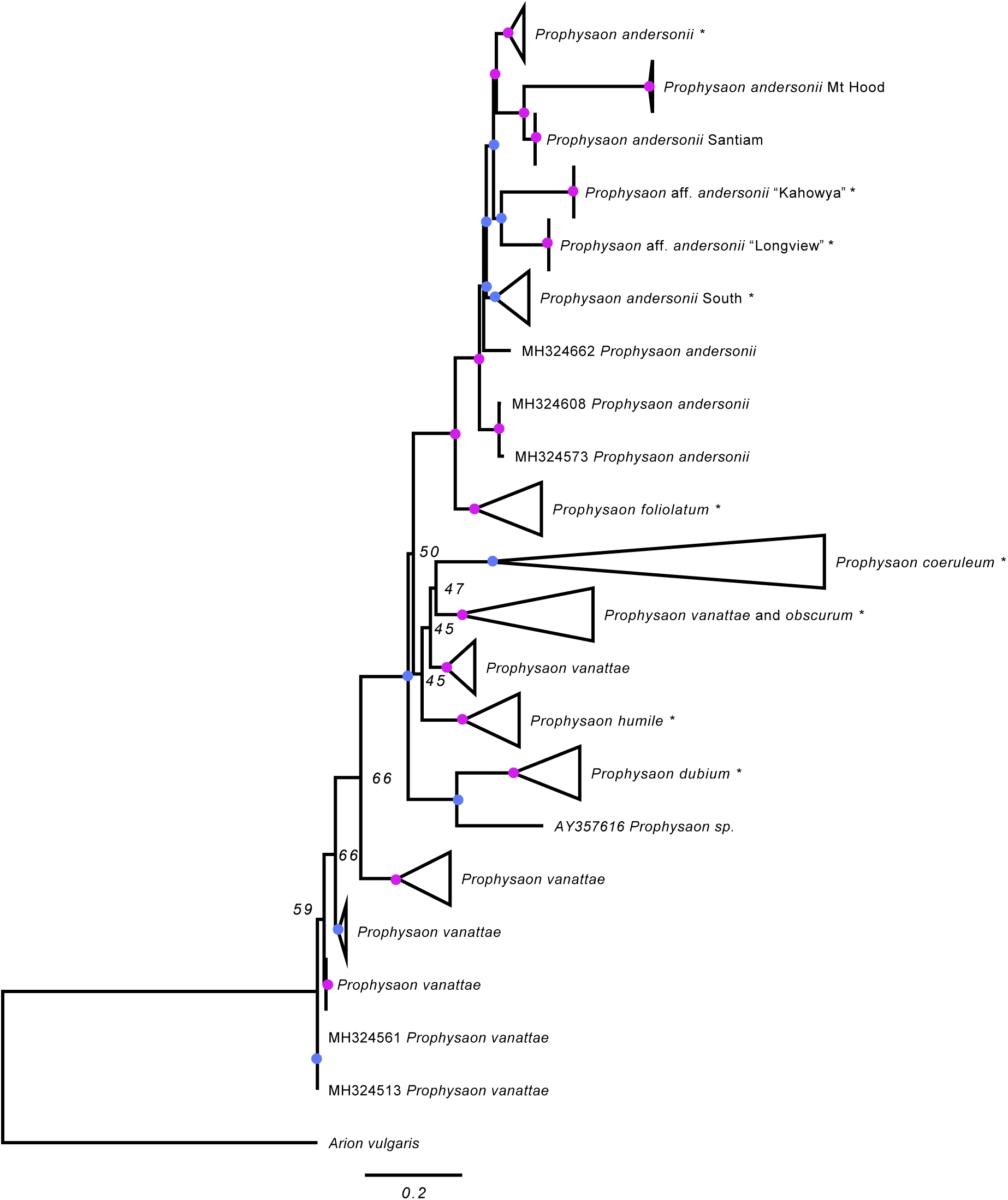
Maximum Likelihood tree inferred for the mitochondrial gene COI. Asterisks indicate clades containing samples from this study. Blue circles indicate nodes with ≥ 75% bootstrap support, while purple circles indicate nodes with ≥ 95% bootstrap support.

### Morphological data

To quantify variation in ecologically relevant morphology, we took scanning electron micrographs of the radulae of *P. andersonii* (n=11)*, P. foliolatum* (n=21)*, P.* aff. *andersonii* “Klahowya” (n=5), and *P.* aff. *andersonii* “Longview” (n=2). The PCA did not show any notable differences between species (Fig. 7). Species was a significant predictor of shape based on the Procrustes ANOVA (p=0.0038). However, after correcting for multiple testing, shape did not differ significantly between any species at an α of 0.01. One of our 2 specimens of *P.* aff. *andersonii* “Longview” had aberrant and poorly mineralized radulae and we suspect it may be a juvenile. Since gastropod radulae can differ substantially in shape, size, and number of teeth through ontogeny within species (Sterki, 1893) (Supporting Fig. S9), we reran our analyses omitting this specimen. In the analysis excluding the apparent juvenile, species was not a significant predictor of shape at an α of 0.01 (p=0.0210).

**Figure 7.**
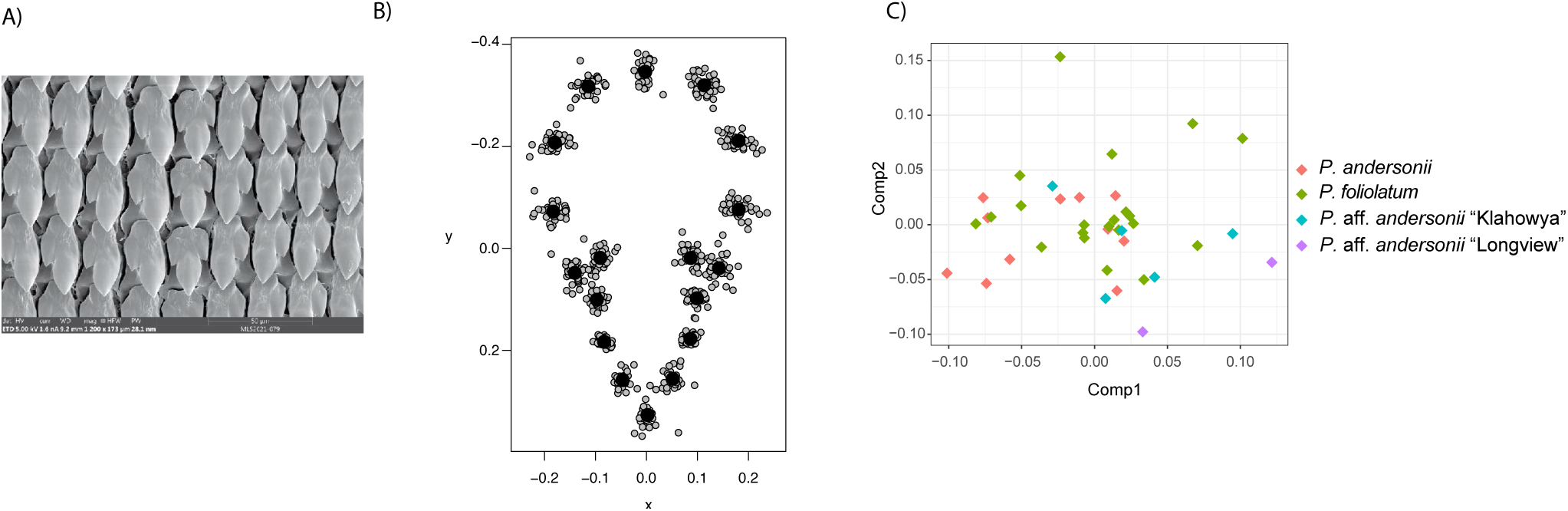
Geometric morphometrics of rachidian shape. A) An exemplar radula from *P. foliolatum* (MLS2021_079). B) The results of GPA on all central rachidia from the *P. andersonii—P. foliolatum* species complex. C) The PCA on rachidian shape, colored by population of origin.

## DISCUSSION

### Phylogeny and subgeneric classification of <u>Prophysaon</u>

For the first time, we were able to infer phylogenetic relationships among species of *Prophysaon* using genomic-scale data. Previous attempts using mitochondrial data (Smith *et al*., 2018) failed to resolve many relationships, particularly those deeper in the tree. Using the transcriptomes collected here, we were able to robustly infer relationships among species.

Despite poor completeness based on BUSCO scores (Supporting Table S1), we were able to assemble a large number of gene families for downstream phylogenetic inference, whether we used all gene families or focused on single-copy genes (Supporting Table S3). Furthermore, the inferred topology was highly stable across datasets and inference methods, and all branches received high bootstrap support and local posterior probabilities. Low concordance factors in parts of the tree likely reflect high levels of ILS due to relatively rapid speciation. Relationships were largely as anticipated. Given their high level of phenotypic similarity, a sister relationship between *P. andersonii* and *P. foliolatum* was unsurprising. That *P. vanattae* was recovered as sister to the other *Mimetarion* sampled, *P. humile,* is consistent with the assertion by Smith et al. that *P. vanattae* and *P. humile* are sister species distributed on either side of the Columbia Basin (2018).

In the trees inferred here, *P. (Prophysaon)—*as it is currently constituted—is paraphyletic, as one of its members, *P. (Prophysaon) coeruleum*, is sister to all other *Prophysaon* species, and *Mimetarion* is a monophyletic clade nested within *P. (Prophysaon)*. However, excluding *P. coeruleum*, the clades *P. (Prophysaon)* sensu stricto and *Mimetarion* are reciprocally monophyletic. The type species of *Prophysaon* is *P. hemphilli* Bland & W.G. Binney, 1874 [“1873”], a junior synonym of *P. andersonii*, and the type species of *Mimetarion* is *P. vanattae*, so these subgeneric names could potentially be conserved by naming a new monotypic subgenus for the *P. coeruleum* clade.

Instead of introducing another name, we hereby synonymize the subgenus *Mimetarion* with nominotypical *Prophysaon* to simplify the nomenclature of the genus, as the original morphological basis for the subgenera no longer holds. Pilsbry based the subgenus *Mimetarion* on reproductive anatomical differences between its species and the rest of the genus *Prophysaon*: in *P. vanattae*, *P. humile*, and *P. fasciatus*, the male reproductive tract is significantly shorter, thinner, and simpler than in the other *Prophysaon* species, which all possess a long, convoluted epiphallus that terminates in a characteristic, massive, muscularized section (Pilsbry, 1948). The fact that the sister species to all other *Prophysaon, P. coeruleum,* possesses the highly specialized epiphallic complex of nominotypical *Prophysaon* suggests that this suite of genital modifications is a synapomorphy of the genus which was subsequently lost in *Mimetarion*.

### Undescribed diversity in the <u>P. andersonii—P.foliolatum</u> species complex

We observed undescribed phenotypic diversity when collecting specimens from the Olympic Peninsula (Klahowya) and near Longview, Washington. These slugs differed noticeably from *P. foliolatum* and *P. andersonii* in their coloration (Fig. 2; Supporting Fig. S3). Structure analyses (Fig. 4) identified these two populations as distinct from *P. andersonii* and *P. foliolatum*, and phylogenetic inference based on transcriptomes placed *P.* aff. *andersonii* “Klahowya” as sister to *P. andersonii* and *P.* aff. *andersonii* “Longview” as sister to *P. foliolatum*. On the other hand, results from the mitochondrial gene COI included broader sampling and placed these two populations as sister to each other and nested within *P. andersonii.* We did not find notable differences in an ecologically relevant trait—radular (i.e. tooth) shape—after removing an outlier from the Longview population. Future research should investigate additional morphological traits from these populations, including reproductive morphology, and sample more broadly from type localities of previously described taxa and other phenotypically distinct populations to better resolve whether these populations represent species-level diversity, or phenotypic variation within *P. andersonii* and *P. foliolatum*.

### The role of introgression in the diversification of <u>Prophysaon</u>

D-statistics support a moderate role of introgression in the history of *Prophysaon*. Not surprisingly, given their phenotypic similarity, our results support introgression between some members of the *P. andersonii*—*P. foliolatum* species complex. Additionally, though, our results support introgression across described subgenera, between *P. vanattae* and *P. humile* and the *P. andersonii*—*P. foliolatum* species complex. Given the high degree of overlap in current species ranges (Supporting Fig. S1), along with potentially reduced ranges and increased overlap during glacial cycles, this is perhaps also not surprising.

### Species of Prophysaon not included in this study

We did not sample 3 species that some authors (e.g., Burke, 2013) treat as valid. All are known only from single lots and have never been recollected. *P. boreale* is only known from its type locality Dyea, Alaska, where we did not collect, and its type series (6 wet specimens, ANSP 87416) has been missing from the Academy of Natural Sciences of Philadelphia Malacology Collection since at least May 2017 (P. Callomon, pers. comm. to NFS, Feb. 2026). *P. fasciatum* is also only known from its type locality, Mendocino Co., California, which was outside of our study region, and its type series is apparently also lost. *P. obscurum* was named as an infrasubspecific color variety of *P. fasciatum* based on a single lot from Chehalis, Washington, and was only nearly a century later elevated to specific rank and removed from the synonymy of the nominal taxon by Branson (1977), but with no discernable justification. Clarifying the taxonomic status of these names and other available names currently in synonymy will be the subject of future revisionary work.

## CONCLUSION

By using transcriptomes, we were able to, for the first time, confidently infer species-level relationships in *Prophysaon*. These results demonstrated that the current morphology-based subgeneric classification is unsupported by molecular data, leading us to synonymize *Mimetarion* with nominotypical *Prophysaon.* We also uncovered previously undescribed genetic and phenotypic diversity within the *P. andersonii—P. foliolatum* species complex. Broader geographic sampling, including from type localities of described species, will facilitate future work clarifying the taxonomy of this group.

## ACKNOWLEDGEMENTS

We would like to thank Karen Smith for assistance with field sampling, and Paul Callomon of the Academy of Natural Sciences of Philadelphia for photographs of and information about the type specimens of *Prophysaon* held by that institution. We would also like to thank Casey Richart for helpful conversations regarding variation within the *P. andersonii—P. foliolatum* species complex and Matthew Hahn for helpful discussions about this work. We used the High-Performance Computing resources available at Indiana University, and thus, this research was supported in part by Lilly Endowment, Inc., through its support for the Indiana University Pervasive Technology Institute.

## AUTHOR CONTRIBUTIONS

Conceptualization, M.L.S.; Data curation, M.L.S., S.M., and N.F.S.; Formal analysis, M.L.S.; Funding acquisition, M.L.S., Methodology, M.L.S. and N.F.S.; Writing—original draft, M.L.S. and N.F.S.; Writing—review & editing, M.L.S. and N.F.S.

## SUPPLEMENTARY DATA

Supplementary data is available online.

## DATA AVAILABILITY

Raw reads are available on the NCBI Short Read Archive (PRJNA1205348). All input files for phylogenetic inference, population genetic inference, and morphological analyses are available on FigShare (doi: 10.6084/m9.figshare.31838968). Vouchers will be deposited at The Ohio State Museum of Biological Diversity upon acceptance.

