## Supporting Information for "Transcriptomes resolve phylogenetic relationships and reveal undescribed diversity in taildropper slugs (Genus *Prophysaon*)"

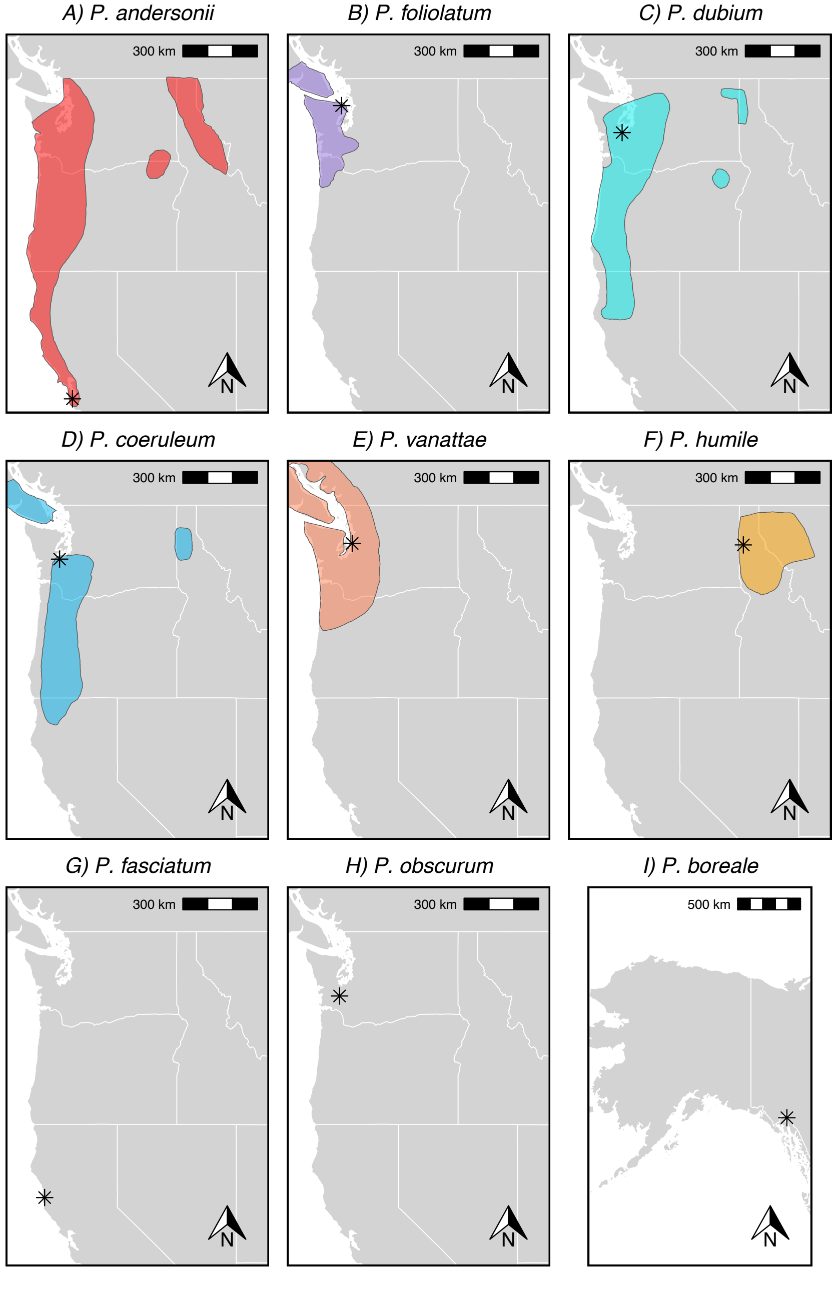


**Supporting Fig. S1.** Distributions and type localities of currently recognized *Prophysaon* species. Subgenus *Prophysaon* includes *P. andersonii* (A), *P. foliolatum* (B), *P. dubium* (C), *P. coeruleum* (D), and *P. boreale* (I). Subgenus *Mimetarion* includes *P. vanattae* (E), *P. humile* (F), *P. obscurum* (H), and *P. fasciatum* (G). The star symbol on each map indicates the type locality for that species. Note that some species are only known from their type localities.

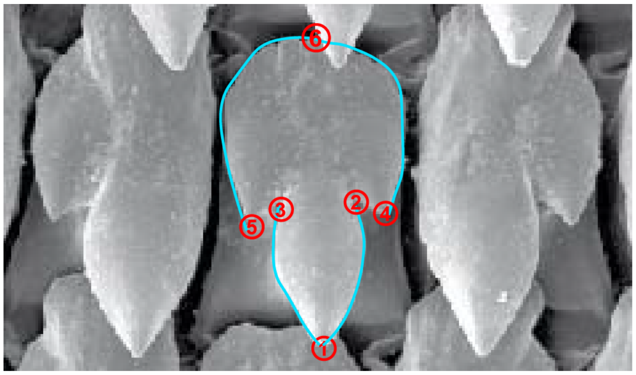


**Supporting Fig. S2.** Landmarks used for geometric morphometric analyses. We focused on the rachidian (central) tooth and placed six landmarks (indicated by circles). We also included four curves (indicated by blue lines).

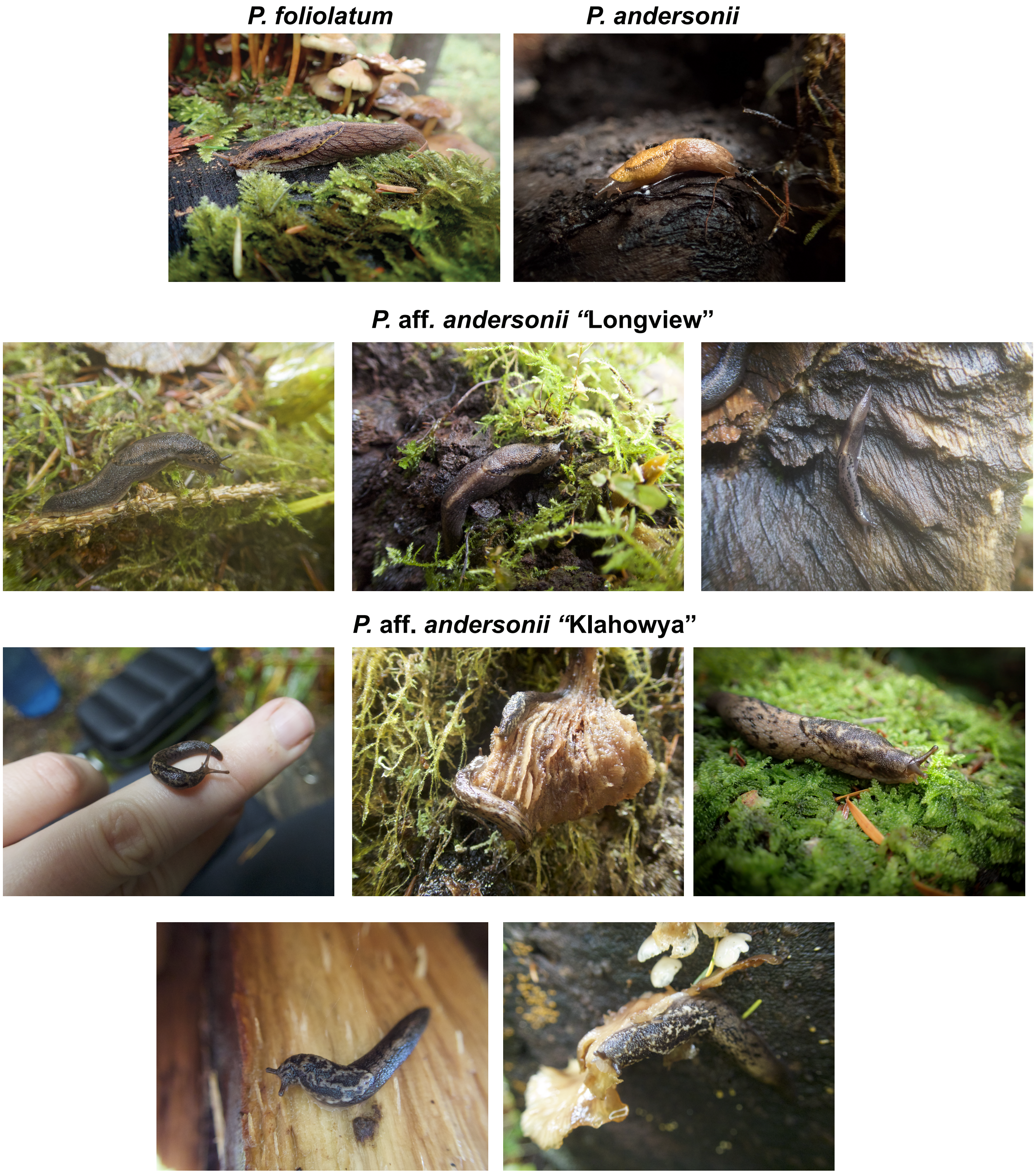


**Supporting Fig. S3.** Additional images of *P. andersonii*, *P. foliolatum*, and phenotypically unique slugs from Longview, Washington and the Klahowya campground in Olympia National Forest. All photos taken by M. Smith.

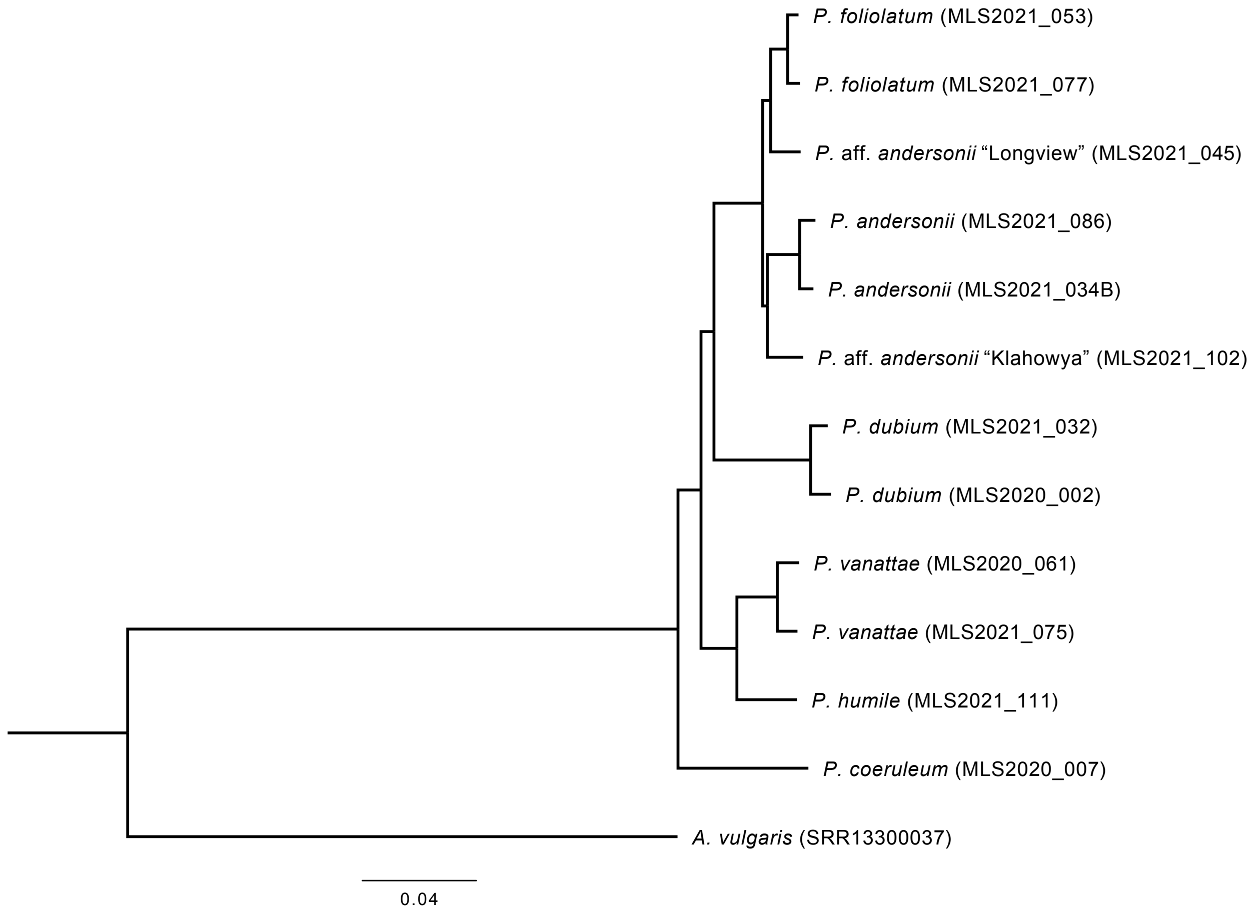


**Supporting Fig. S4.** Maximum Likelihood tree inferred from single-copy genes in IQ-Tree. Maximum Likelihood trees inferred in IQ-Tree were topologically identical for all datasets analyzed. Bootstrap support values in IQ-Tree were 100 across all nodes and datasets.

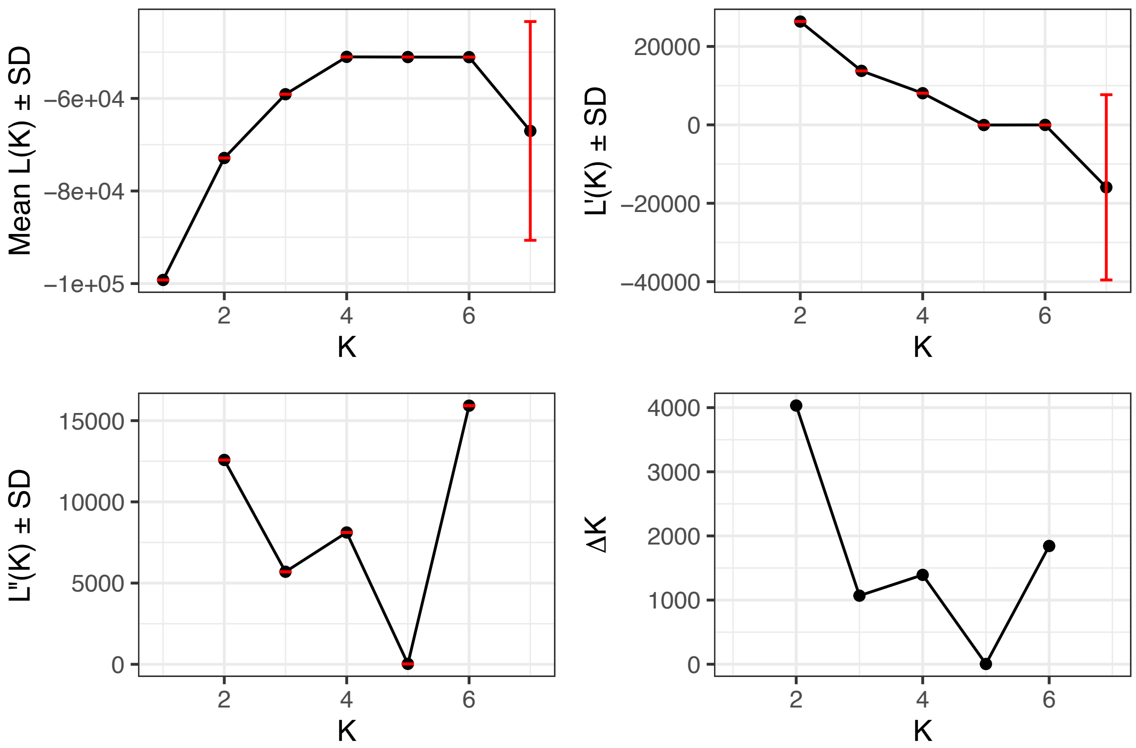


**Supporting Fig. S5.** L(K) is the posterior probability of the data for a given K. L\`(K) is the mean difference between successive likelihood values of K. L\`\`(K) is the absolute value of the difference between successive values of L\`(K). ΔK is the mean absolute value of L\`\`(K) averaged over runs and divided by the standard deviation of L(K).

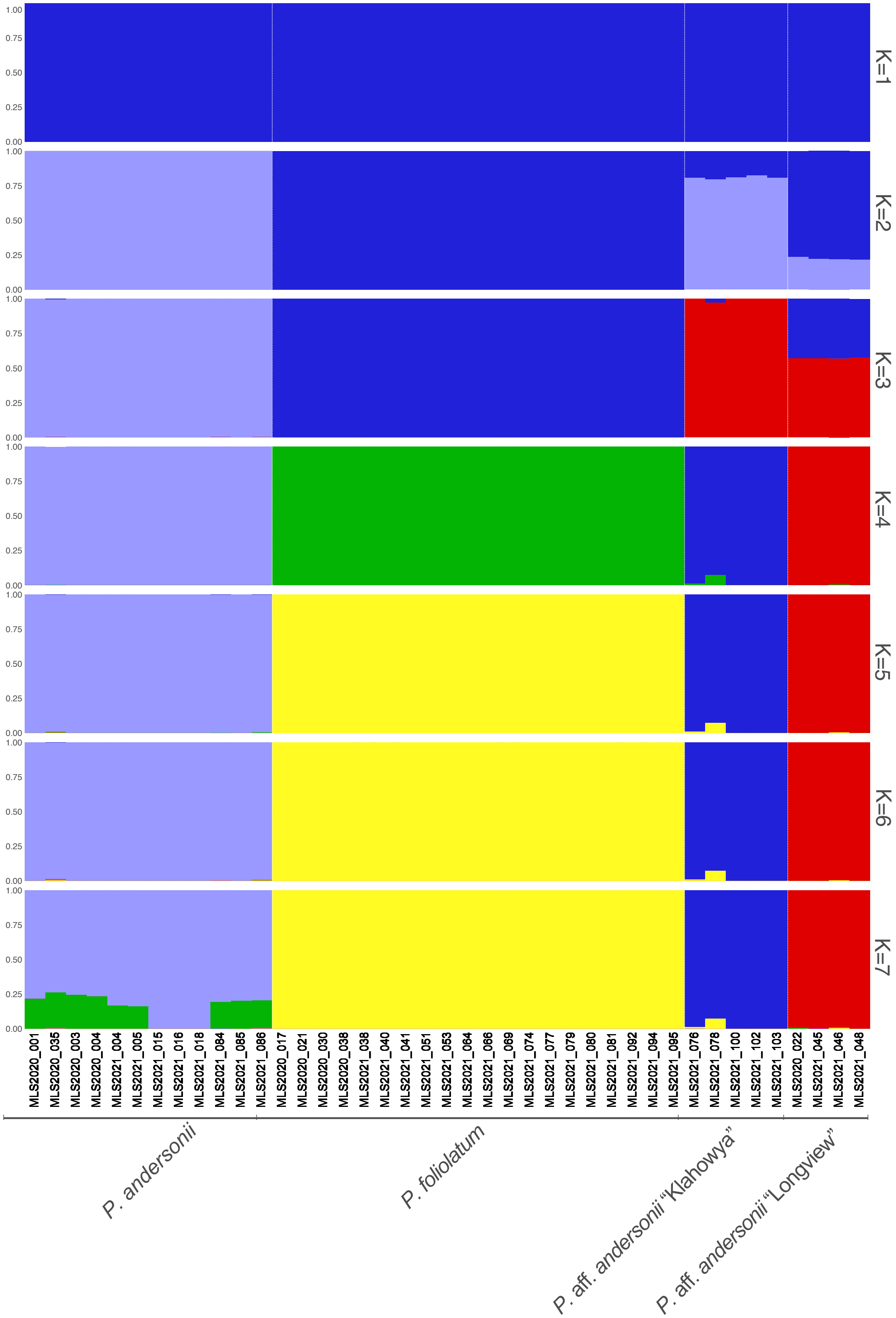


**Supporting Fig. S6.** Population assignments from Structure for K-1 through K=7.

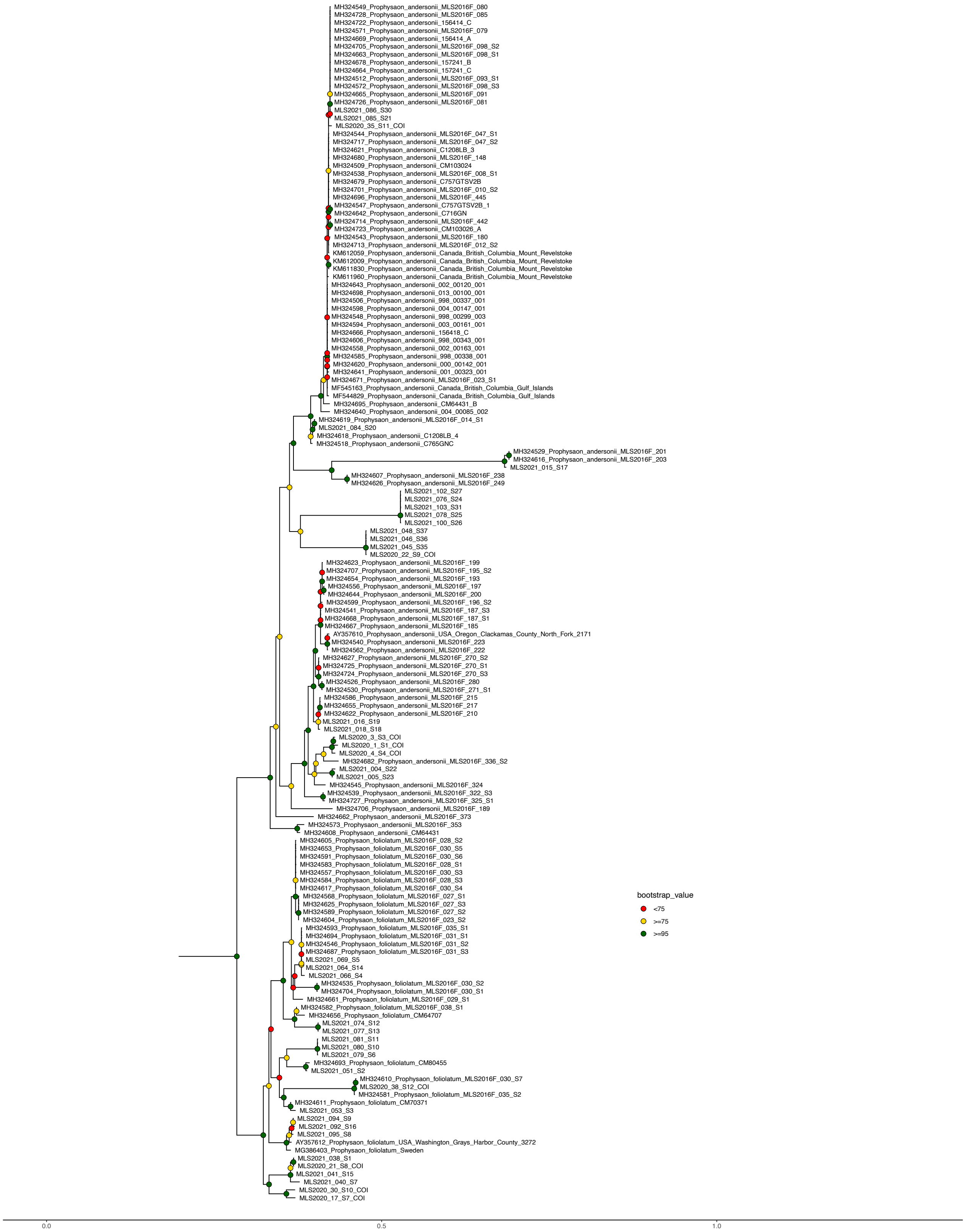


**Supporting Fig. S7.** COI tree with full labels for the *P. andersonii-P.foliolatum* species complex.

**Supporting Fig. S8.** COI tree with full labels for all taxa except the *P. andersonii-P.foliolatum* species complex.

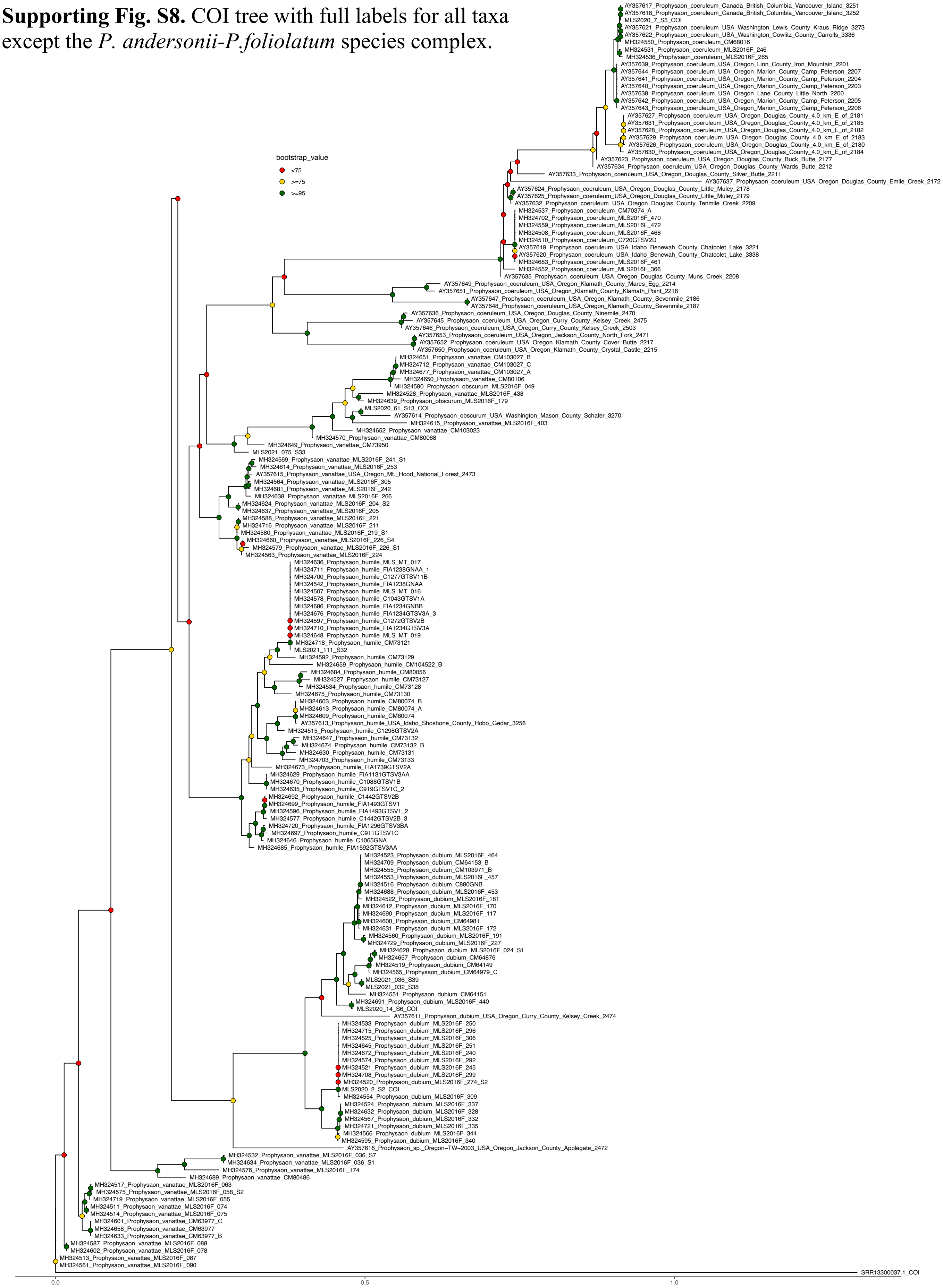

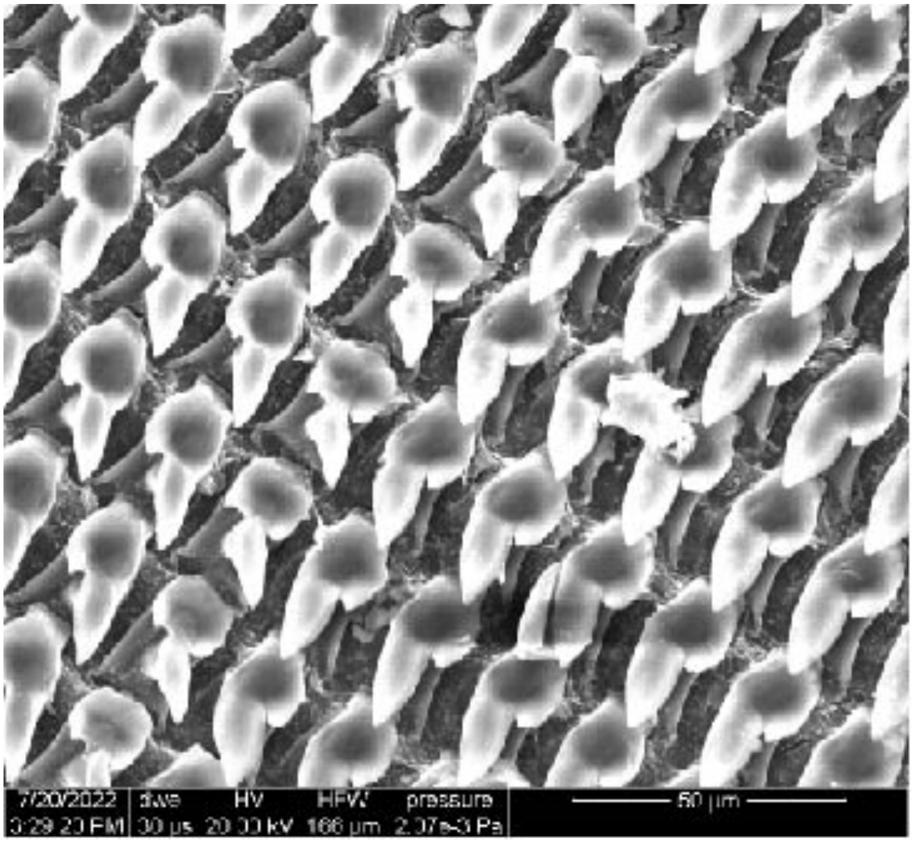


**Supporting Fig. S9.** Scanning electron micrograph of the aberrant radula of the apparently juvenile “Longview” specimen (MLS2021_046. Note the “bent” appearance of the rachidian (central) teeth in comparison to Supporting Fig. S2 and Fig. 7 in the main text.

**Supporting Table S1: Transcriptome Assemblies.** Trinity and BUSCO statistics for transcriptome assemblies. For source of tissue, all indicates that foot, albumin gland, digestive gland, epiphallus, and mantle covering were used.

| Sample | Species | Total trinity genes | Contig N50 | Median contig length | Complete (BUSCO) | Single-Copy (BUSCO) | Duplicated (BUSCO) | Fragmented (BUSCO) | Missing (BUSCO) | Source of Tissue | Latitude | Longitude |
|---|---|---|---|---|---|---|---|---|---|---|---|---|
| MLS2020_001 | *P. andersonii* | 45741 | 944 | 428 | 1023 | 923 | 100 | 119 | 4153 | all | 44.7678833 | -122.6265 |
| MLS2020_002 | *P. dubium* | 52264 | 839 | 424 | 1449 | 1329 | 120 | 138 | 3708 | all | 44.7678833 | -122.6265 |
| MLS2020_003 | *P. andersonii* | 36271 | 859 | 437 | 824 | 747 | 77 | 90 | 4381 | all | 44.7678833 | -122.6265 |
| MLS2020_004 | *P. andersonii* | 62547 | 975 | 443 | 1558 | 1432 | 126 | 156 | 3581 | all | 44.7678833 | -122.6265 |
| MLS2020_007 | *P. coeruleum* | 72403 | 1041 | 479 | 2360 | 2144 | 216 | 167 | 2768 | foot, albumin gland, epiphallus, mantle covering | 45.38245 | -122.037367 |
| MLS2020_014 | *P. dubium* | 51438 | 794 | 408 | 1293 | 1180 | 113 | 152 | 3850 | all | 45.3807667 | -122.02995 |
| MLS2020_017 | *P. foliolatum* | 62057 | 919 | 423 | 1707 | 1567 | 140 | 148 | 3440 | all | 45.7804833 | -122.35175 |
| MLS2020_021 | *P. foliolatum* | 32509 | 859 | 424 | 850 | 783 | 67 | 98 | 4347 | albumin gland | 45.7959667 | -123.458633 |
| MLS2020_022 | *P.* aff. *andersonii* “Longview” | 66454 | 933 | 426 | 1681 | 1523 | 158 | 170 | 3444 | all | 45.9565333 | -123.578683 |
| MLS2020_030 | *P. foliolatum* | 42212 | 829 | 410 | 904 | 820 | 84 | 101 | 4290 | all | 46.1772333 | -123.501633 |
| MLS2020_035 | *P. andersonii* | 23222 | 790 | 411 | 565 | 509 | 56 | 65 | 4665 | all | 46.0681333 | -122.8531 |
| MLS2020_038 | *P. foliolatum* | 38881 | 905 | 434 | 1001 | 917 | 84 | 104 | 4190 | albumin gland | 46.9239167 | -123.061383 |
| MLS2020_061 | *P. vanattae* | 68880 | 886 | 433 | 1604 | 1444 | 160 | 177 | 3514 | all | 47.4924833 | -123.209833 |
| MLS2021_004 | *P. andersonii* | 73421 | 938 | 413 | 1827 | 1662 | 165 | 171 | 3297 | albumin gland and ephiphallus | 44.3940333 | -122.24545 |
| MLS2021_005 | *P. andersonii* | 74427 | 943 | 398 | 1998 | 1819 | 179 | 198 | 3099 | albumin gland and ephiphallus | 44.3940333 | -122.24545 |
| MLS2021_015 | *P. andersonii* | 62522 | 1037 | 419 | 2015 | 1855 | 160 | 154 | 3126 | albumin gland and ephiphallus | 45.2985833 | -122.360883 |
| MLS2021_016 | *P. andersonii* | 59027 | 944 | 399 | 1605 | 1466 | 139 | 171 | 3519 | albumin gland and ephiphallus | 45.2985833 | -122.360883 |
| MLS2021_018 | *P. andersonii* | 48547 | 871 | 390 | 1301 | 1195 | 106 | 153 | 3841 | albumin gland and ephiphallus | 45.2985833 | -122.360883 |

| | | | | | | | | | | | | |
|---|---|---|---|---|---|---|---|---|---|---|---|---|
| MLS2021_032 | *P. dubium* | 128830 | 851 | 383 | 2891 | 2535 | 356 | 250 | 2154 | albumin gland and ephiphallus | 45.78015 | -123.3516 |
| MLS2021_036 | *P. dubium* | 87270 | 810 | 379 | 2239 | 2031 | 208 | 231 | 2825 | albumin gland and ephiphallus | 45.78015 | -123.3516 |
| MLS2021_038 | *P. foliolatum* | 43093 | 840 | 381 | 992 | 897 | 95 | 132 | 4171 | albumin gland and ephiphallus | 45.7960833 | -123.458867 |
| MLS2021_040 | *P. foliolatum* | 31841 | 803 | 381 | 809 | 730 | 79 | 94 | 4392 | albumin gland and ephiphallus | 45.7960833 | -123.458867 |
| MLS2021_041 | *P. foliolatum* | 58024 | 553 | 295 | 790 | 731 | 59 | 85 | 4420 | albumin gland and ephiphallus | 45.7960833 | -123.458867 |
| MLS2021_045 | *P.* aff. *andersonii* “Longview” | 94452 | 849 | 379 | 1940 | 1745 | 195 | 213 | 3142 | albumin gland and ephiphallus | 45.9565 | -123.578783 |
| MLS2021_046 | *P.* aff. *andersonii* “Longview” | 54951 | 800 | 377 | 970 | 885 | 85 | 112 | 4213 | albumin gland and ephiphallus | 45.9565 | -123.578783 |
| MLS2021_048 | *P.* aff. *andersonii* “Longview” | 75141 | 900 | 398 | 1500 | 1364 | 136 | 160 | 3635 | albumin gland and ephiphallus | 45.9565 | -123.578783 |
| MLS2021_051 | *P. foliolatum* | 65192 | 922 | 388 | 1550 | 1439 | 111 | 153 | 3592 | albumin gland and ephiphallus | 46.19285 | -123.164383 |
| MLS2021_053 | *P. foliolatum* | 80132 | 1008 | 414 | 2206 | 2025 | 181 | 192 | 2897 | albumin gland and ephiphallus | 46.19285 | -123.164383 |
| MLS2021_064 | *P. foliolatum* | 45527 | 951 | 391 | 1420 | 1302 | 118 | 125 | 3750 | albumin gland and ephiphallus | 47.0981833 | -123.464083 |
| MLS2021_066 | *P. foliolatum* | 50282 | 687 | 342 | 852 | 769 | 83 | 120 | 4323 | albumin gland and ephiphallus | 47.0981833 | -123.464083 |
| MLS2021_069 | *P. foliolatum* | 74515 | 750 | 356 | 1400 | 1272 | 128 | 174 | 3721 | albumin gland and ephiphallus | 47.0981833 | -123.464083 |
| MLS2021_074 | *P. foliolatum* | 40897 | 785 | 380 | 995 | 908 | 87 | 126 | 4174 | albumin gland and ephiphallus | 48.0639833 | -124.109983 |
| MLS2021_075 | *P. vanattae* | 106782 | 834 | 384 | 2347 | 2118 | 229 | 218 | 2730 | albumin gland and ephiphallus | 48.0639833 | -124.109983 |
| MLS2021_076 | *P.* aff. *andersonii* “Klahowya” | 68861 | 694 | 363 | 1241 | 1122 | 119 | 137 | 3917 | albumin gland and ephiphallus | 48.0639833 | -124.109983 |
| MLS2021_077 | *P. foliolatum* | 69898 | 738 | 362 | 1274 | 1160 | 114 | 147 | 3874 | albumin gland and ephiphallus | 48.0639833 | -124.109983 |

| | | | | | | | | | | | | |
|---|---|---|---|---|---|---|---|---|---|---|---|---|
| MLS2021_078 | *P.* aff. *andersonii* “Klahowya” | 49793 | 628 | 344 | 764 | 686 | 78 | 103 | 4428 | albumin gland and ephiphallus | 48.0639833 | -124.109983 |
| MLS2021_079 | *P. foliolatum* | 26119 | 684 | 355 | 554 | 495 | 59 | 77 | 4664 | albumin gland and ephiphallus | 48.1168833 | -124.2348 |
| MLS2021_080 | *P. foliolatum* | 61303 | 686 | 354 | 889 | 829 | 60 | 87 | 4319 | albumin gland and ephiphallus | 48.1168833 | -124.2348 |
| MLS2021_081 | *P. foliolatum* | 72243 | 753 | 364 | 1242 | 1138 | 104 | 130 | 3923 | albumin gland and ephiphallus | 48.1168833 | -124.2348 |
| MLS2021_084 | *P. andersonii* | 49349 | 843 | 369 | 982 | 886 | 96 | 137 | 4176 | albumin gland and ephiphallus | 47.5957167 | -123.122367 |
| MLS2021_085 | *P. andersonii* | 75982 | 813 | 369 | 1504 | 1344 | 160 | 187 | 3604 | albumin gland and ephiphallus | 47.5957167 | -123.122367 |
| MLS2021_086 | *P. andersonii* | 90664 | 944 | 392 | 2035 | 1812 | 223 | 214 | 3046 | albumin gland and ephiphallus | 47.5957167 | -123.122367 |
| MLS2021_092 | *P. foliolatum* | 71755 | 547 | 306 | 790 | 730 | 60 | 90 | 4415 | albumin gland and ephiphallus | 47.4623667 | -123.8577 |
| MLS2021_094 | *P. foliolatum* | 31842 | 736 | 367 | 675 | 620 | 55 | 70 | 4550 | albumin gland and ephiphallus | 47.4623667 | -123.8577 |
| MLS2021_095 | *P. foliolatum* | 33576 | 787 | 367 | 820 | 744 | 76 | 116 | 4359 | albumin gland and ephiphallus | 47.4623667 | -123.8577 |
| MLS2021_100 | *P.* aff. *andersonii* “Klahowya” | 101242 | 735 | 342 | 1866 | 1674 | 192 | 206 | 3223 | albumin gland and ephiphallus | 48.0639833 | -124.109983 |
| MLS2021_102 | *P.* aff. *andersonii* “Klahowya” | 103201 | 899 | 392 | 2773 | 2482 | 291 | 208 | 2314 | albumin gland and ephiphallus | 48.0639833 | -124.109983 |
| MLS2021_103 | *P.* aff. *andersonii* “Klahowya” | 68196 | 794 | 381 | 1607 | 1458 | 149 | 159 | 3529 | albumin gland and ephiphallus | 48.0639833 | -124.109983 |
| MLS2021_111 | *P. humile* | 61103 | 898 | 401 | 1745 | 1566 | 179 | 179 | 3371 | albumin gland and ephiphallus | 48.19045 | -116.158033 |

**Supporting Table S2.** Specimens used for geometric morphometric analyses.

| Specimen | Species | Latitude | Longitude |
|---|---|---|---|
| MLS2017F_036 | *P. andersonii* | 44.7804 | -122.50927 |
| MLS2017F_089 | *P. andersonii* | 45.32889 | -122.17973 |
| MLS2017F_130 | *P. foliolatum* | 46.92681 | -123.06163 |
| MLS2017F_131 | *P. foliolatum* | 46.87975 | -123.22681 |
| MLS2017F_139 | *P. foliolatum* | 47.09779 | -123.46554 |
| MLS2017F_144 | *P. foliolatum* | 46.99654 | -123.59076 |
| MLS2020_003 | *P. andersonii* | 44.7678833 | -122.6265 |
| MLS2020_004 | *P. andersonii* | 44.7678833 | -122.6265 |
| MLS2020_017 | *P. foliolatum* | 45.7804833 | -122.35175 |
| MLS2020_021 | *P. foliolatum* | 45.7959667 | -123.458633 |
| MLS2020_022 | *P.* aff. *andersonii* "Longview" | 45.9565333 | -123.578683 |
| MLS2020_030 | *P. foliolatum* | 46.1772333 | -123.501633 |
| MLS2020_035 | *P. andersonii* | 46.0681333 | -122.8531 |
| MLS2020_038 | *P. foliolatum* | 46.9239167 | -123.061383 |
| MLS2021_004 | *P. andersonii* | 44.3940333 | -122.24545 |
| MLS2021_005 | *P. andersonii* | 44.3940333 | -122.24545 |
| MLS2021_015 | *P. andersonii* | 45.2985833 | -122.360883 |
| MLS2021_018 | *P. andersonii* | 45.2985833 | -122.360883 |
| MLS2021_038 | *P. foliolatum* | 45.7960833 | -123.458867 |
| MLS2021_040 | *P. foliolatum* | 45.7960833 | -123.458867 |
| MLS2021_041 | *P. foliolatum* | 45.7960833 | -123.458867 |
| MLS2021_046 | *P.* aff. *andersonii* "Longview" | 45.9565 | -123.578783 |
| MLS2021_051 | *P. foliolatum* | 46.19285 | -123.164383 |
| MLS2021_064 | *P. foliolatum* | 47.0981833 | -123.464083 |
| MLS2021_069 | *P. foliolatum* | 47.0981833 | -123.464083 |
| MLS2021_074 | *P. foliolatum* | 48.0639833 | -124.109983 |
| MLS2021_076 | *P.* aff. *andersonii* "Klahowya" | 48.0639833 | -124.109983 |
| MLS2021_077 | *P. foliolatum* | 48.0639833 | -124.109983 |
| MLS2021_086 | *P. andersonii* | 47.5957167 | -123.122367 |
| MLS2021_092 | *P. foliolatum* | 47.4623667 | -123.8577 |
| MLS2021_094 | *P. foliolatum* | 47.4623667 | -123.8577 |
| MLS2021_095 | *P. foliolatum* | 47.4623667 | -123.8577 |
| MLS2021_102 | *P.* aff. *andersonii* "Klahowya" | 48.0639833 | -124.109983 |
| MLS2021_078 | *P.* aff. *andersonii* "Klahowya" | 48.0639833 | -124.109983 |
| MLS2021_079 | *P. foliolatum* | 48.1168833 | -124.2348 |
| MLS2021_080 | *P. foliolatum* | 48.1168833 | -124.2348 |
| MLS2021_084 | *P. andersonii* | 47.5957167 | -123.122367 |
| MLS2021_100 | *P.* aff. *andersonii* "Klahowya" | 48.0639833 | -124.109983 |
| MLS2021_103 | *P.* aff. *andersonii* "Klahowya" | 48.0639833 | -124.109983 |

**Supporting Table S3.** The number of gene trees in each phylogenetic dataset. The all genes dataset includes all gene families, whether or not they contain paralogs. The single-copy orthologs datasets includes only those gene families present in a single-copy in any sampled species. The Maximum Inclusion, Monophyletic Outgroups, and DISCO datasets include gene families after they were trimmed using branch-cutting techniques.

| **Dataset** | **Genes** |
|---|---|
| All Genes | 8081 |
| Single-copy Orthologs | 7315 |
| Maximum Inclusion | 8441 |
| Monophyletic Outgroups | 7731 |
| DISCO | 8457 |
